# Genetic exchange shapes ultra-small Patescibacteria metabolic capacities in the terrestrial subsurface

**DOI:** 10.1101/2022.10.05.510940

**Authors:** Emilie Gios, Olivia E. Mosley, Nobuto Takeuchi, Kim M. Handley

## Abstract

Bacterial genomes are highly dynamic entities, mostly due to the extent of horizontal gene transfer (HGT) occurring in these organisms. HGT is thought to be the main driver of genetic variation and adaptation to local environment in bacteria. However, little is known about the modalities of HGT within natural microbial communities, especially the implications of genetic exchange for streamlined microorganisms such as Patescibacteria (Candidate Phyla Radiation). We searched for evidence of genetic exchange in 125 Patescibacteria genomes recovered from aquifer environments and detected the presence of hundreds of genomic islands, individually transferred genes and prophage combined, with up to 29% of genome length attributed to HGT. Results show that most individual gene transfer events occurred between Patescibacteria, but donors were also phylogenetically diverse groundwater microorganisms. Using gene donor-recipient information, we identified one potential host (Omnitrophota) of the ultra-small bacteria, and confirmed this by matching relative abundance patterns across 16 groundwater samples. A wide variety of metabolic functions were introduced in Patescibacteria genomes by HGT including transcription, translation and DNA replication, recombination and repair. This study illustrates the evolutionarily dynamic nature of Patescibacteria genomes despite the constraints of streamlining, and that HGT in these organisms is also mediated via viral infection.

## INTRODUCTION

The horizontal transfer of genetic material (or horizontal gene transfer; HGT), involving acquisition of exogenous DNA, is thought to be a key factor in the evolution of prokaryotes, especially in bacteria (Ochman et al., 2000). HGT accounts for a significant fraction of gene diversity and genomic fluidity in these organisms, leading to a variety of genome structures and physiological diversity (Soucy et al., 2015). This evolutionary process is important for niche adaptation, and is often driven by mobile genetic elements (MGEs) such as genomic islands (GIs), transposons and phage. Many routes exist for gene transfer: uptake of environmental DNA molecules (transformation), exchange between two cellular organisms via a physical bond (conjugation), or delivery through phage infection (transduction).

Recent metagenomic studies of subsurface ecosystems reveal a large group of unusual bacteria referred to as the Patescibacteria phylum (equating to the Candidate Phyla Radiation; CPR), which has been suggested to comprise up to 26% of all currently known bacterial diversity (Parks et al., 2017; Schulz et al., 2017). The group constitutes taxa consistently harbouring small cell and genome sizes (Brown et al., 2015). Implications of reduced size include limited metabolic functions, such as the lack of known amino acid and nucleotide biosynthesis pathways (Castelle et al., 2018). These observations led to the prediction that most ultra-small Patescibacteria have a symbiotic lifestyle (Brown et al., 2015), and some were confirmed to be episymbionts based on cultivation and microscopy studies (He et al., 2021). Close physical associations with other organisms due to metabolic dependencies, along with the long residence time of groundwater (Griebler & Lueders, 2009), and the importance of the biofilms in aquifers (Flynn et al., 2008), could favour genetic exchange in subsurface ultra-small microbial communities.

Despite proximity to other groundwater bacteria, extensive gene loss in Patescibacteria evolution could result in selective pressures against the acquisition and retention of exogenous DNA that are stronger than in non-streamlined genomes. Another potential barrier to HGT lies in the fact that genome reduction in obligate intracellular symbionts has often been associated with loss of functions involved in DNA recombination and repair (Moran & Bennett, 2014), which could limit integration of foreign DNA. Previous studies have, however, shown not only that a significant fraction of Patescibacteria organisms still possesses genes required for homologous recombination (Castelle et al., 2018), but that they are naturally competent for uptake of eDNA molecules, mediated by DNA intake pumps (via comEC and pili complexes; Castelle et al., 2017, 2018; Nelson & Stegen, 2015).

Investigating the mobility of adaptive genes via HGT could provide important information about the evolutionary strategy of these ultra-small organisms, including aspects governing putative host-symbiont interactions. Implications of HGT include altered interaction modalities with host populations, as described in *Vibrio fischeri*. Adding a single gene to the *V. fischeri* genome is sufficient to alter its host range in squids (Mandel et al., 2009). Additionally, acquisition of MGEs has been shown to constitute evidence of symbiotic relationships between HGT recipient and donor, such as extensive gene transfer from *Wolbachia* endosymbionts to their insect hosts during genome reduction (Dunning Hotopp et al., 2007), and putative acquisition of host-derived ankyrin repeat proteins by bacterial sponge symbionts to inhibit phagocytosis (Nguyen et al., 2014). How potentially adaptive genes aid niche adaptation in ultra-small prokaryotes and fine-tune metabolic interactions with their predicted ‘basibiont’ prokaryotic host cells remains largely unknown.

To understand the role of HGT in the evolution of complex, natural groundwater communities, and particularly streamlined Patescibacteria, we analyzed 396 metagenome-assembled genomes (MAGs), including 132 Patescibacteria genomes. We investigated genomic evidence for, and the frequency of, individual horizontal gene transfer events (defined here as HT genes) in the MAGs, and compared these to acquisitions of larger MGEs such as GIs and prophage. Results provide evidence for many HGT events among Patescibacteria, and exchanges with other prokaryotes in groundwater. We report that metabolic functions horizontally acquired by Patescibacteria taxa appear to be biased towards features distinct from the general groundwater communities.

## MATERIALS AND METHODS

### Sample collection

Groundwater samples were collected from 8 wells from 2 sand/gravel aquifers in Canterbury, New Zealand, as described by (Mosley et al., 2022). Briefly, wells were purged (×3–5 bore volumes), then 3–90 L of groundwater was filtered per sample. Biomass was captured on 0.22 μm mixed cellulose ester (MCE) filters, after passing through a 1.2 μm MCE pre-filter, using a 142 mm filter holder (Merck Millipore Ltd., Cork, Ireland). Samples were preserved in RNAlater (ThermoFisher Scientific, Waltham, MA, USA), transported on dry ice, and stored at −80°C.

After collecting a groundwater sample (as above), a second sample per well was collected following low frequency sonication (2.43 kW for 2 minutes) to induce biofilm (and sediment) detachment from the surrounding aquifer, using a custom sonication device (Close et al., 2020). 0.5–15 L of sediment (and hence biomass) enriched groundwater was filtered as above.

### Nucleic acid extraction

To remove RNAlater, filters were centrifuged (2500 g for 5 min) and washed with nuclease-free phosphate-buffered saline (pH 7.4, ThermoFisher Scientific, Waltham, MA, USA). DNA was extracted using the DNeasy PowerSoil Pro Kit (Qiagen, Valencia, CA, USA) and 0.25-0.9 g of whole filter per reaction. Samples were extracted in replicate (1–47 reactions/sample). Replicates were pooled and concentrated using a sodium acetate/ethanol precipitation with glycogen (0.1 μg/μL final concentration, Roche, Basel, Switzerland). High molecular weight DNA was confirmed via gel electrophoresis. DNA was quantified using Qubit fluorometry (ThermoFisher Scientific), and quality was checked using a NanoPhotometer (Implen, Munich, Germany). Extractions yielded 70–864 ng of DNA (8.7 ng for gwj02) for metagenomics.

### Metagenome sequencing

Whole genome shotgun sequencing was undertaken on the 16 Canterbury samples (gwj01-16). DNA libraries were prepared using the Illumina TruSeq Nano DNA Kit with ∼550 bp inserts, by Otago Genomics (University of Otago, Dunedin, NZ). The low-yield gwj02 sample was prepared with the ThruPLEX DNA-Seq Kit (Takara Bio USA, Inc., Mountain View, CA, USA). Sequences (2×250 bp) were generated using the Illumina HiSeq 2500 V4 platform. Adapter sequences were removed using Cutadapt v2.10 (Martin, 2011), and reads were quality trimmed using sickle v01.33 (parameters -q 30 −l 80). Read quality was checked using FastQC v0.11.7 (https://www.bioinformatics.babraham.ac.uk/projects/fastqc/).

### Metagenome assembly and binning

The 16 metagenomes were individually assembled using SPAdes v3.11.1 (Bankevich et al., 2012) (parameters: --meta, -k 43,55,77,99,121). Metagenomes from the same well also were co-assembled (same parameters). A dereplicated set of metagenome-assembled genomes (MAGs) (99% average nucleotide identity, ANI, dereplication threshold) was generated as described previously (Tee et al., 2020), and refined according to bin_detangling script (https://github.com/dwwaite/bin_detangling).

For ultra-small prokaryotes, genome completeness and contamination were assessed based on the presence of 51 bacterial single-copy genes (SCGs) for Dependentiae, 43 SCGs for Patescibacteria and 38 archaeal SCGs for DPANN (Anantharaman et al., 2016). For other prokaryotes, completeness was estimated using CheckM v1.0.13 (Parks et al., 2015). To determine coverage, reads were mapped onto the de-replicated MAGs using Bowtie (Langmead et al., 2009). Sample-specific genome relative abundance was calculated by normalizing based on the highest read count between samples (Probst et al., 2018).

### Gene prediction and functional annotation

Protein coding sequences were predicted using Prodigal v2.6.3 (Hyatt et al., 2010). Genes were annotated using USEARCH (Edgar, 2010) against UniRef100 (Suzek et al., 2015), Uniprot (The UniProt Consortium, 2017) and KEGG (Kanehisa et al., 2016) databases (e-value ≤0.001 and identity ≥50%), and HMMs against PFAM (Sonnhammer et al., 1998) and TIGRFAM (Haft et al., 2003) databases using HMMER v3.1b2 (Potter et al., 2018) (e-value ≤0.001). Additionally, functional annotation was carried out using eggNOG-mapper v2 (Huerta-Cepas et al., 2017) against the eggNOG v5.0 database (Huerta-Cepas et al., 2019), using default parameters.

### Genome taxonomy

Taxonomic classification of MAGs was performed using the Genome Taxonomy Database Toolkit (GTDB-Tk, v0.2.1) (Chaumeil et al., 2020) with database release 89 (Parks et al., 2020). Maximum likelihood trees were constructed using 120 bacterial and 122 archaeal concatenated marker genes with IQ-TREE v1.6.12 (L.-T. Nguyen et al., 2015) using 100 bootstraps, and ModelFinder (Kalyaanamoorthy et al., 2017) best-fit model LG+F+G4 for bacteria and LG+F+I+G4 for archaea. Trees were visualized and annotated with iTOL (Letunic & Bork, 2007). MAGs (≥80% complete) were compared to GTDB representative genomes (downloaded 20-Jan-2020) by calculating Average Amino-acid Identities (AAI) for blastp matches sharing ≥30% identity over ≥70% of alignment length.

### Identification of horizontal gene transfer

Individual HT events among MAGs were identified using MetaCHIP v1.10.0 (Song et al., 2019) at phylum, class, order, family and genus levels. False-positive HT genes (potentially introduced via assembly contamination) were filtered out by removing HT genes marked by MetaCHIP as “full-length match” (where the match represents a large proportion of contig) or “end match” (HT genes located at the end of contig) as described by Dong et al. (2021). Information about HGT events involving members of the Patescibacteria phylum were retrieved and visualised using the Circlize package in R (Gu et al., 2014).

For genomic island prediction, contigs were concatenated into contiguous sequences and uploaded to IslandViewer4 web server (Bertelli et al., 2017) for genomic island prediction. GIs that were predicted by at least one method, and that did not overlap two concatenated contigs, were considered for the analysis. Genes located on GIs were compared to community gene content using blastn.

Prophage integrated in the Patescibacteria genomes were identified using VIBRANT 1.2.1 (Kieft et al., 2020). Whenever a predicted viral sequence was overlapping a predicted GI, CheckV quality assessment information was used and only “High-quality” genomes were retained as prophage (Nayfach et al., 2021).

### Phylogenetic distance

Average Amino-acid Identities (AAI) between MAGs (≥80% complete) involved in HGT events were determined via blastp matches sharing ≥30% identity over ≥50% of query and database alignment length.

### Orthology analysis

To investigate the Patescibacteria pangenome, we performed an orthologous clustering of 118,635 protein-coding sequences encoded by the recovered MAGs. Pairwise all-vs-all blastp searches were conducted using the following cut-offs: ≥30% nucleotide identity over ≥50% of query sequence length, e-value ≤0.001. BLAST table outputs were used as input for orthogroup clustering using Orthofinder v2.3.1 (Emms & Kelly, 2019) with -b option and default parameters.

## RESULTS AND DISCUSSION

### Prevalence of HGT events in Patescibacteria genomes

Using the community-level HGT detection tool MetaCHIP (Song et al., 2019), we investigated the potential horizontal transfer of individual genes between aquifer microorganisms. A total of 1,487 transfers were identified among the 396 groundwater MAGs (Table S1), including 1,407 unique HT genes. Taxonomic breakdown of detected HT events revealed that over half of the Patescibacteria genomes experienced HT with another genome in the dataset, regardless of their taxonomy or estimated genome size (Fig. 1a,b). One hundred and twenty-three (9% of) horizontally acquired genes were identified in 68 Patescibacteria organisms, representing 1.0 ±1.2 HT (average ± standard deviation) genes received per genome and 1.1 ± 1.3 HT genes per Mbp (Fig. 1b). In comparison, genomes of other members of the community contained on average 3.5 ± 4.8 horizontally transferred genes per genome, but a comparable 1.4 ± 2.1 per Mbp. The comparison of HT events per Mbp indicates that the rate of exchange of individual genes is mostly independent of genome size (i.e. larger genomes do not receive proportionally more genes). This is supported by significant positive correlations between HT events per genome and estimated genome size (both donor and recipient HT events, Fig. 2), and is consistent with previous observations of HT acquired genes in prokaryotic genomes (Garcia-Vallvé et al., 2000), and of transfers occurring between distantly related organisms (i.e. cross-order transfers, >85% of the total HT events detected here) (Cordero & Hogeweg, 2009).

**Figure 1.**
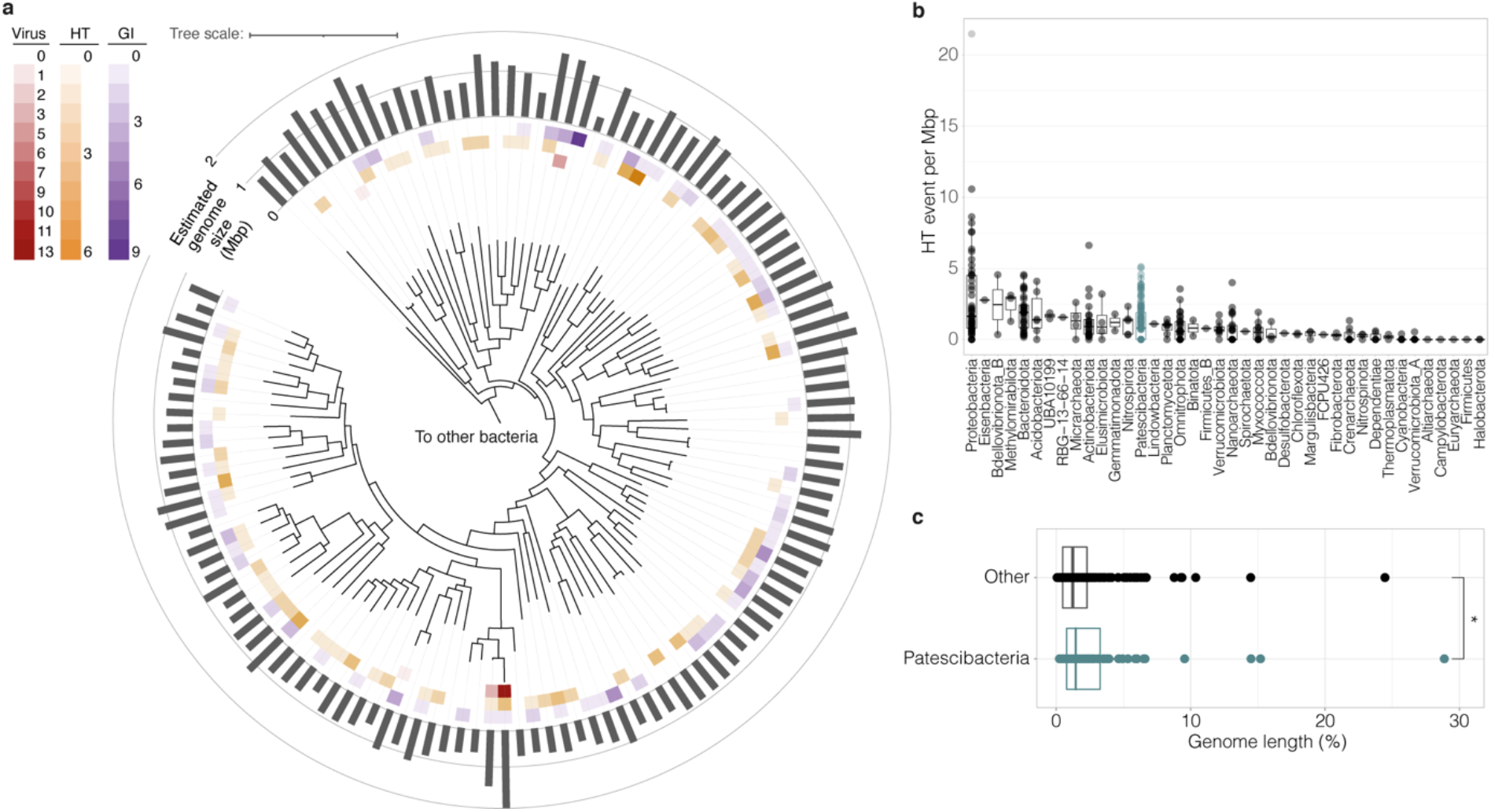
Prevalence of HGT in Patescibacteria and the wider groundwater community. (a) Maximum likelihood phylogenetic trees of 132 Patescibacteria MAGs recovered in this study. Trees are based on 120 concatenated bacterial marker gene alignment from GTDB-Tk. Rings from the center: number of prophage; number of HT events; number of GIs; estimated genome size (Mbp). The scale bar indicates the number of substitutions per site. (b) Number of recipient HT events between all groundwater MAGs per phylum normalized by genome size. Patescibacteria phylum is indicated in teal. The center line of each boxplot represents the median; the top and bottom lines are the first and third quartiles, respectively; and the whiskers show 1.5× the interquartile range. (c) Proportion of each genome occupied by acquired GIs and prophage sequences, comparing Patescibacteria (teal) and the rest of the community (black). Significant differences were assessed for each group using Wilcoxon Signed Rank (*: p < 0.05).

**Figure 2.**
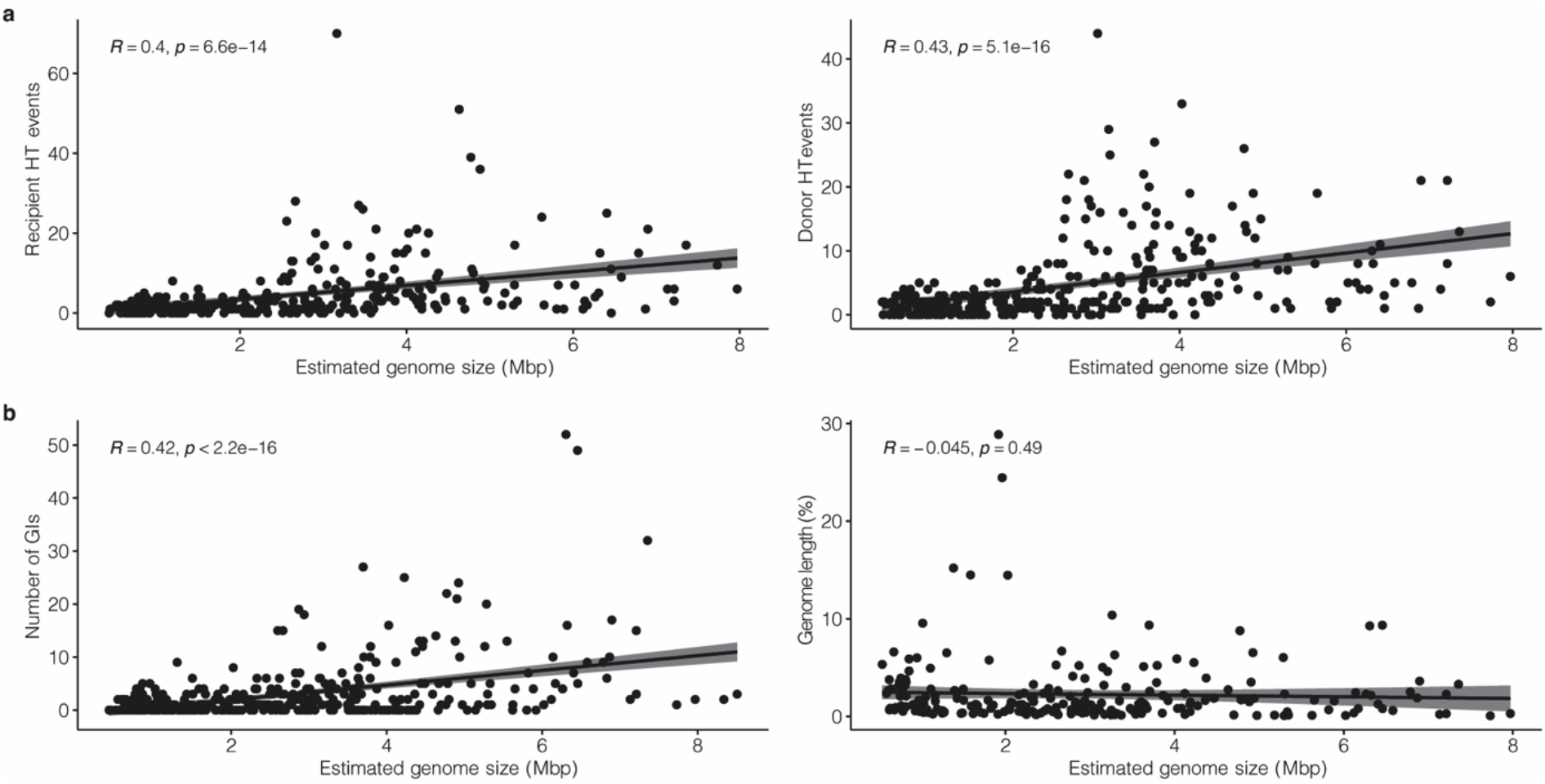
Correlations between the estimated genome size of groundwater-derived MAGs and HGT. (a) Estimated genome size compared with recipient HT events (left) and donor HT events (right). (b) Estimated genome size compared with the number of GIs detected (left), and the proportion of genome length occupied by GI and prophage sequences (right).

When searching for large MGEs, we found 120 GIs in 63 Patescibacteria MAGs, ranging from 2.7 to 45 kbp long, with 40 GIs over 10 kb long (average length 17.4 ± 0.7 kb) (Table S2). Additionally, 24 putative prophage genomes (30% of those overall) were found to be concentrated in 5 Patescibacteria MAGs – with the proportion of prophage in Patescibacteria being comparable to the proportion of Patescibacteria MAGs in the community (33%). This frequency of prophage in ultra-small bacterial genomes is surprising, as prophage frequency has been shown to increase with genome size (Touchon et al., 2016). Accordingly, our analysis shows that, as also observed for HT genes, the frequency of acquisition of longer complex DNA sequences, constituting GIs, increased with the genome size of groundwater-derived MAGs (Fig. 2b).

When considering the groundwater community overall, the proportion of genome length occupied by GIs and prophages, combined, remained constant regardless of recipient genome size (Fig. 2b), and despite GI and prophage sequences being similar in length between ultra-small and other microorganisms (Fig. 3, Table S2). This is consistent with observations, by Ochman et al. (2000), that the proportion of horizontally transferred genes is independent of the amount of protein-coding sequences in complete prokaryotic genomes. However, when comparing the two community fractions, results here show the proportion of genome length occupied by putative GIs and prophage was slightly, but significantly higher in Patescibacteria than the rest of the communities – on average 3.0 ± 4.4% versus 2.0 ± 2.8%, respectively (Fig. 1c).

**Figure 3.**
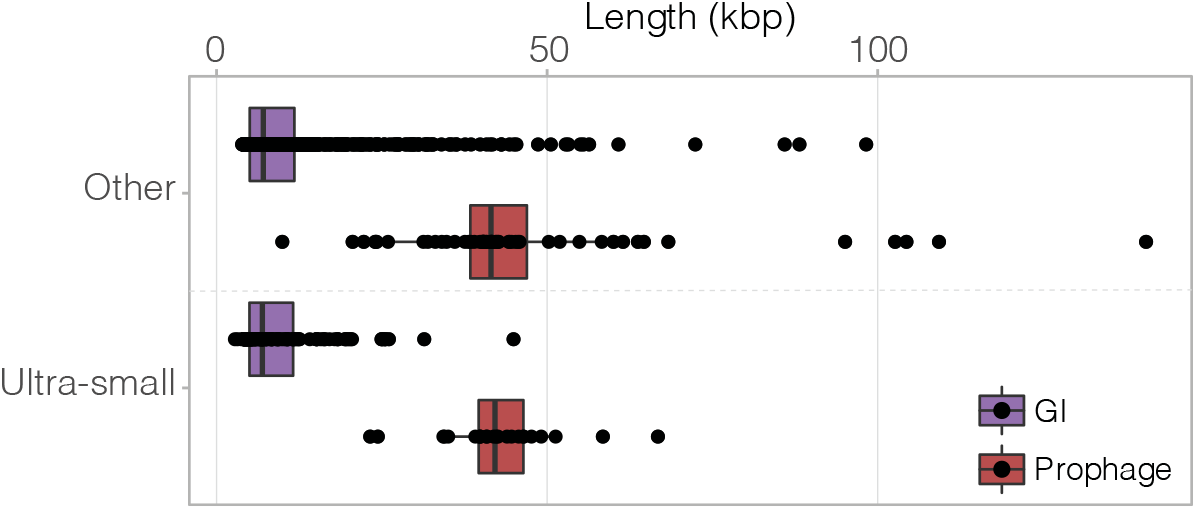
Length distribution of GIs and prophage in ultra-small MAGs and other groundwater microbial MAGs. The center line of each boxplot represents the median; the top and bottom lines are the first and third quartiles, respectively; and the whiskers show 1.5× the interquartile range. No significant differences were detected in GI and prophage sequence length between ultra-small and other groundwater microorganisms (Wilcoxon Signed Rank, p>0.05).

The extent of gene transfer events detected here is comparable to those detected in a variety of other phylogenetically diverse microbial communities, e.g. 90.4% of groundwater MAGs containing detectable HGT events, compared to 78.8% in cheese-associated communities, and 89.5% in prokaryotic isolate genomes (Bonham et al., 2017; Ochman et al., 2000). However, the extent of genetic exchange observed in the groundwater microbial community is likely underestimated. Ancient transfer events will escape detection due to robust integration within the recipient genome via modification of GC content and tetranucleotide frequencies (Lawrence & Ochman, 1997). Moreover, while MetaCHIP has proven efficient in predicting HGT donors/recipients relationships within microbial communities, donor taxa might not have been captured in the MAG dataset.

### HGT origins in Patescibacteria taxa

Most HT events detected occurred between members of the Patescibacteria (Fig. 4a), with 104 of the 123 HT genes received by Patescibacteria originating from counterparts. This observation is consistent with research, which demonstrates that HGT is most prevalent among closely related organisms (Bonham et al., 2017), and that intra-species HGT occurs at far greater frequency (Majewski et al., 2000). HGT was likely also facilitated by the spatial proximity of microorganisms (Gogarten et al., 2002), in particular by microbial cohorts in groundwater communities (Hug et al., 2015). This includes cohorts formed by ultra-small prokaryotes, where individuals spatially co-occur as described previously in subsurface and soil ecosystems (Geesink et al., 2020; Herrmann et al., 2019; Nicolas et al., 2021).

**Figure 4.**
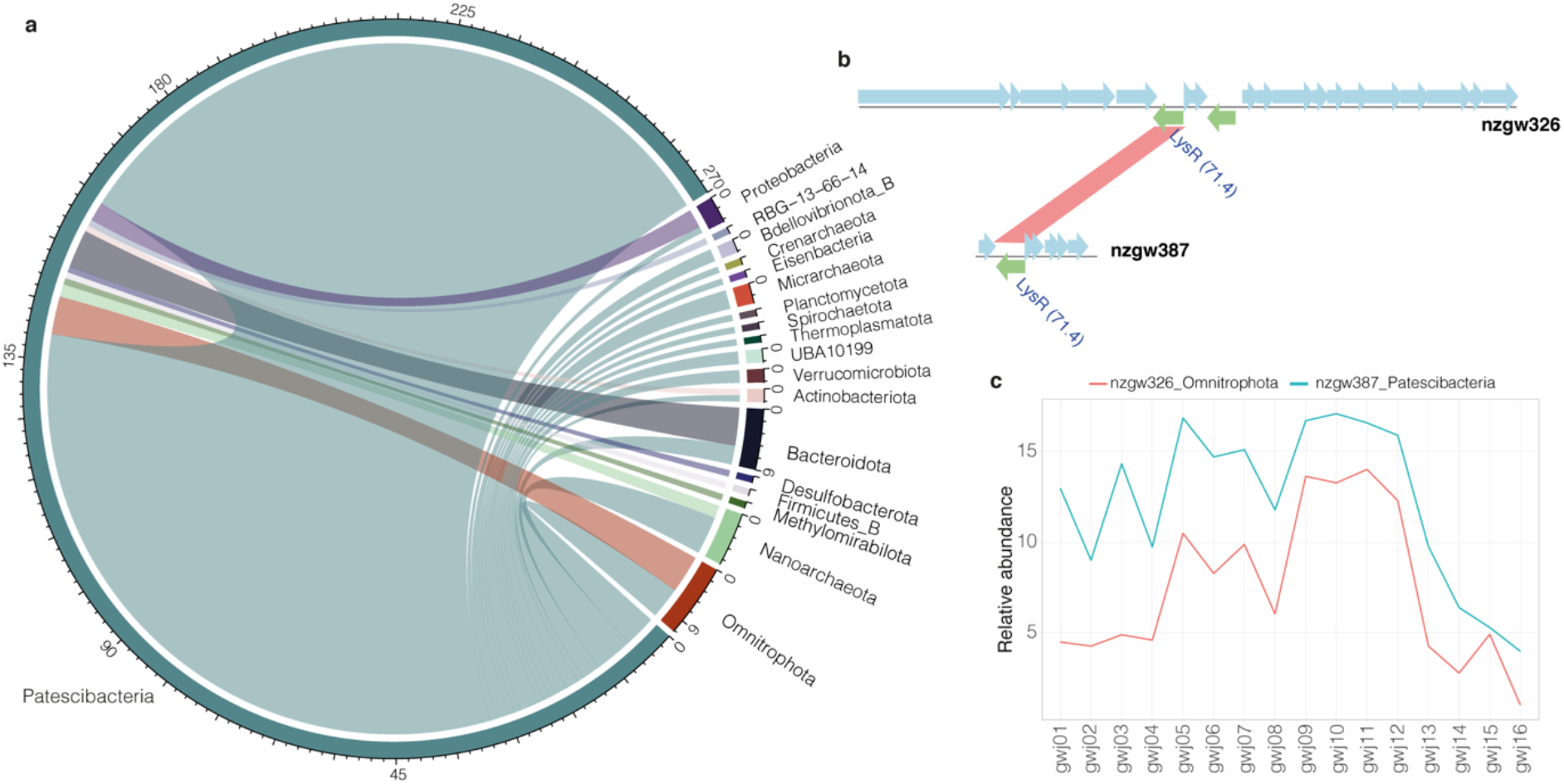
Origins of HT genes in Patescibacteria taxa. (a) Gene flow among phyla donating to Patescibacteria taxa. Scale represents number of HT events and inner bands show donor/recipient relationship (coloured according to donor organism). (b) Example of HT gene transferred from Omnitrophota nzgw326 (upper plot) to Patescibacteria nzgw387 (lower plot). Genes encoded on the forward strand are displayed in blue, and genes coded on the reverse strand are displayed in green. (c) Genome relative abundance profiles of these 2 organisms across 16 groundwater samples.

Although a smaller number of MAGs were recovered from Omnitrophota and Bacteroidota, organisms in these phyla shared genes more frequently with Patescibacteria (3.5x and 3x more, respectively) than, for example, Proteobacteria, which was the second most populous phylum (Tables 1, S3). Frequent sharing suggests potential close relationships between these organisms and ultra-small prokaryotes (i.e. host/symbiont). The former two bacterial groups have been identified previously as putative HGT donors to Patescibacteria in seafloor environments (Dong et al., 2021). Moreover, Omnitrophota and Bacteroidota organisms were found to consistently co-occur with the ultra-small bacteria in aquifers (Chaudhari et al., 2021).

**Table 1.** Frequency of HT events to Patescibacteria at phylum level. Note the ratio of donor HGT events to number of MAGs is biases in donor phyla with small MAG sample sizes (Methylomirabilota, Firmicutes_B, Desulfobacterota, RBG-13-66-14).

| Donor phylum | Donor HGT events | Reconstructed donor MAGs | Ratio donor HGT events / number of MAGs |
| --- | --- | --- | --- |
| Patescibacteria | 109 | 132 | 0.83 |
| Omnitrophota | 6 | 28 | 0.21 |
| Bacteroidota | 6 | 33 | 0.18 |
| Nanoarchaeota | 2 | 23 | 0.09 |
| Proteobacteria | 3 | 51 | 0.06 |
| Actinobacteriota | 1 | 24 | 0.04 |
| Methylomirabilota | 1 | 3 | 0.33 |
| Firmicutes_B | 1 | 1 | 1.00 |
| Desulfobacterota | 1 | 1 | 1.00 |
| RBG-13-66-14 | 1 | 1 | 1.00 |

We observed further evidence for recent putative horizontal transfers to groundwater Patescibacteria from patescibacterial counterparts and other taxa based on positive correlations between the relative abundances of HGT pairs across sites (29 out of 131 pairs involving Patescibacteria as donor or recipient with Pearson’s correlation coefficient >0.5, p <0.05, Table S4). For example, Patescibacteria nzgw387 received a copy of the *lysR* gene from Omnitrophota nzgw326 (Fig. 4b), a highly conserved transcriptional regulator involved in numerous cellular functions (Maddocks & Oyston, 2008), and is shown to co-occur with its donor, Omnitrophota nzgw326, in groundwater (Fig. 4c).

A BLAST analysis of GI genes against the gene content of whole groundwater communities showed that most of the genes located on GIs could not be matched to any organisms within the communities (91.9% of GI genes; Table S5). Nonetheless, we found that 7% of GI genes were related to those in Patescibacteria other than the one carrying the GI. Interestingly, the content of one 4,250 bp long GI in Patescibacteria nzgw468 (74% genome completeness, no contamination) was highly similar to genes in the archaeon Halobacterota nzgw24 (99% completeness, 1.3% contaminated), indicating a potential recent transfer event between the two distantly related organisms. None of the 11 genes located on the GI could be functionally characterized (as for a large proportion of genes in Patescibacteria genomes), but were between 85 and 100% identical in amino acid composition to the ones of nzgw24. While inter-domain transfers do occur between Archaea and Bacteria, they are less frequent than bacterial-domain confined HGT events (Nelson et al., 1999, López-García et al., 2015). Gene exchange between Patescibacteria and their archaeal ultra-small counterparts (DPANN) has been reported (León-Zayas et al., 2017), particularly in the context of membrane-associated protein encoding genes potentially involved in host-symbiont interactions (Jaffe et al., 2016). Accordingly, we detected 9 HT events from Patescibacteria to DPANN archaea (Micrarchaeota and Nanoarchaeota; function of transferred genes discussed below). A further, 2 genes (one ABC-transporter and one gene related to the BRO family, of which the function remains unknown) were transferred from Nanoarchaeota to Patescibacteria (Table S6).

### Metabolic functions encoded in horizontally acquired regions

Patescibacteria, or CPR bacteria, have reduced genomes and limited biosynthetic capacities (Castelle et al., 2018). The metabolic function of horizontally transferred genes that are retained in Patescibacteria genomes could provide clues about the evolution of dependencies in these ultra-small prokaryotes regarding their symbiotic interactions with hosts. Although a substantial portion of genes located in horizontally acquired genomic regions could not be assigned any function (61.6% of all genes for Patescibacteria, 38.9% for other taxa), genes located in those regions in Patescibacteria genomes that were able to be annotated are involved in a wide range of metabolic pathways (Fig. 5), and include a large number of genes involved in translation and transcription. Horizontally acquired translation and transcription genes predominantly comprised ribosomal proteins (small and large subunits), elongation factor *tuf*, which plays a central role in protein synthesis, and chaperonin *groS* (Table S6). These results reflect trends identified when considering the most frequently transferred genes to other groundwater prokaryotes, which notably included the *rpsL* gene encoding ribosomal protein S12 and transcription initiation factor *infA*, alongside translation elongation factor *tuf*. Kanhere and colleagues described similar findings (Kanhere & Vingron, 2009). By inferring HGT events based on evolutionary distances within orthologous protein families (using the COG database), authors found that genes exchanged among bacteria were primarily involved in translation (Kanhere & Vingron, 2009).

**Figure 5.**
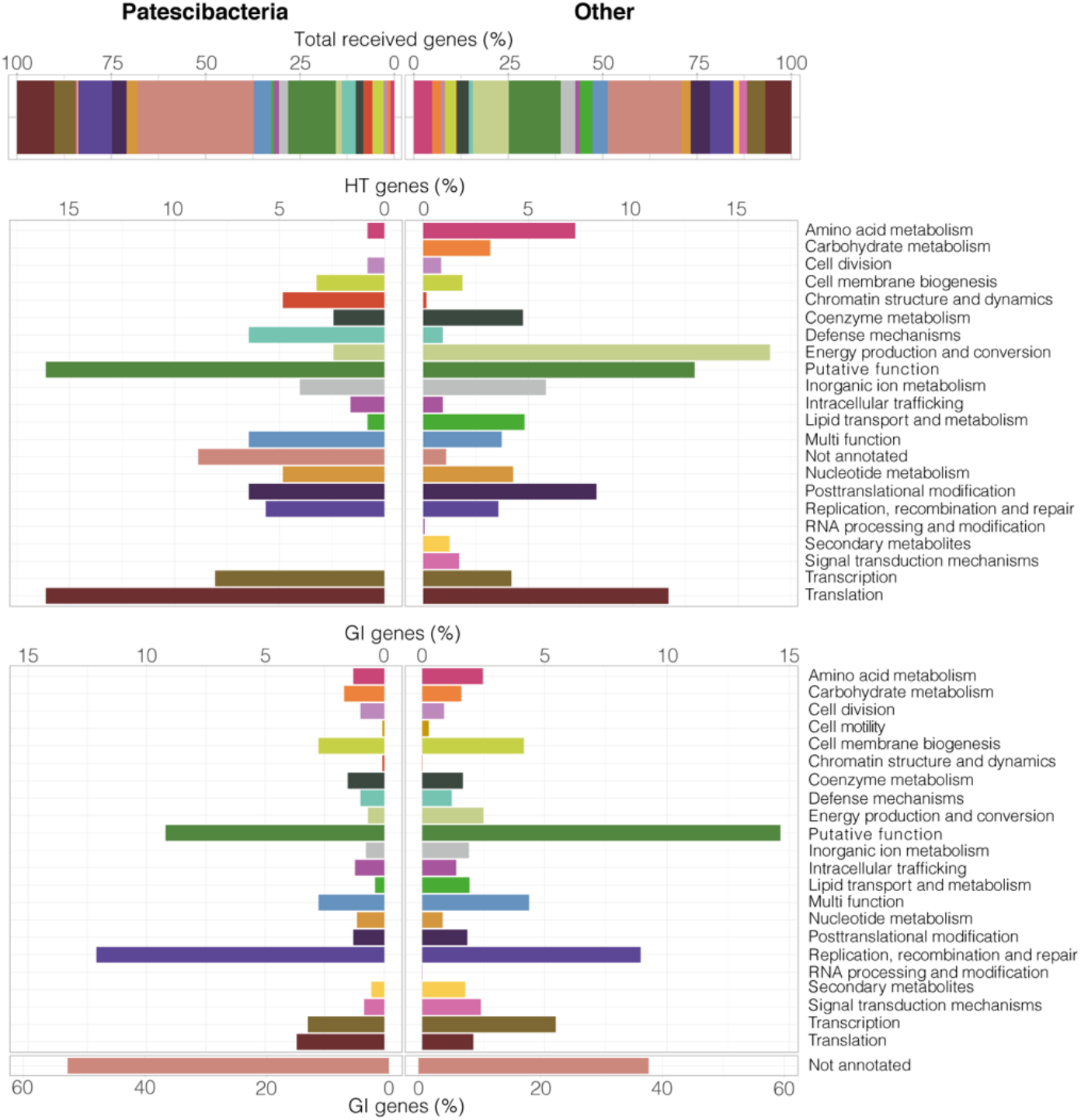
Metabolic functions of genes received through horizontal transfer. Metabolic functions acquired through HGT (HT and GI genes) by Patescibacteria (124 HT genes + 120 GIs received) and other groundwater microorganisms (1,283 HT genes + 926 GIs received) from any other member of the groundwater community (top). Breakdown of metabolic functions of HT genes (middle) and genes located on GIs (bottom).

HT genes in Patescibacteria were found to be highly enriched, compared to other phyla, in genes assigned to the EggNOG “Defense mechanisms” category (Fig. 5). These included genes (e.g. *macB* and *msbA*) encoding ABC-type systems for multidrug and antimicrobial peptide transport. Membrane-located multidrug transporters have been identified previously in relatives of ultra-small bacterium *Babela massiliensis* (Dependentiae phylum, previously Candidate division TM6). Their specialization is thought to be important at the host-symbiont interface (Jaffe et al., 2016). There is growing evidence that drug transporters are required for survival of intracellular pathogenic bacteria in host cells (Du et al., 2015). Genes assigned to chromatin structure were also more prevalent in the HT gene pool in Patescibacteria. Among these genes were six SWIB/MDM2 domain containing proteins, which are chromatin-remodelling proteins involved in transcriptional activation (Tang et al., 2010), and were almost exclusively transferred between Patescibacteria (Table S6). DNA packaging in the form of chromatin is well documented in Eukaryotes and Archaea, but remains poorly understood in Bacteria (Shen & Landick, 2019). Whether ultra-small bacteria use chromatin-remodelling proteins to manipulate their own genomic DNA structure, or for the modification of eukaryotic/archaeal symbiotic host DNA is unclear.

In contrast, other groundwater taxa more frequently acquired genes involved in energy production (respiratory chain) and amino acid biosynthesis, which Patescibacteria are known to consistently lack (Castelle et al., 2018). This could further indicate that ultra-small bacteria must scavenge resources such as protons and amino acids from the host or surrounding environment. A higher proportion of genes involved in secondary metabolite pathways and signal transduction mechanisms was also detected in other taxa (426 HT and GI genes in 105 ‘other’ MAGs versus 18 genes in 15 ultra-small MAGs) (Table S6). Secondary metabolites are considered luxury metabolism (Demain & Fang, 2000) and can constitute public goods (e.g. siderophores) (Weigert & Kümmerli, 2017). As such, biosynthetic gene clusters encoding for these metabolites are uncommon in streamlined genomes (Konstantinidis & Tiedje, 2004; Sharrar et al., 2020), which have reduced auxiliary functions (Giovannoni et al., 2014), and are more likely to depend on the metabolites produced by other organisms (Morris et al., 2012). Similarly, loss of signal transduction mechanisms, which facilitate environmental monitoring, is a characteristic feature of streamlined bacterial genomes, including those of bacteria with non-intracellular lifestyles (e.g. bacterioplankton) (Gabaldón & Huynen, 2007; Swan et al., 2013).

Although it has been shown that gene function impacts HGT overall, studies assessing functional categories of horizontally transferred genes have reported diverging results. The ‘complexity hypothesis’ suggests that genes involved in cellular housekeeping are more frequently subjected to horizontal transfer, while informational genes (involved in transcription, translation, and related processes are more resistant to transfers (Jain et al., 1999). Nakamura et al. (2004) found that horizontally transferred genes are biased towards cell surface, DNA binding and pathogenicity related functions. Conversely, other studies have found that the functional category of a gene family is an insignificant factor in determining HGT (Choi & Kim, 2007; Cohen et al., 2011). While we found that the genes localized in GIs harbour similar metabolic functions overall between Patescibacteria and the rest of the groundwater community, functions of individual genes acquired by the ultra-small bacteria through horizontal transfer tend to differ (Fig. 5). Results obtained here therefore suggest lifestyle of the recipient impacts the traits of genes transferred and also retained.

Biases towards horizontally acquired features distinct from the general groundwater communities by Patescibacteria may be explained by constraints on streamlined genomes. Selection of distinct features could occur by two mechanisms: (i) retention of horizontally transferred genes and concentration due to selective deletion of other less desirable features, and (ii) selective retention of acquired genes with desirable features. The caveat being that the distribution of functions among the hypothetical coding DNA sequence (CDS) fraction in Patescibacteria is unknown.

We could question whether the length of transferred DNA is an important factor affecting integration and retainment of transferred genes and related metabolic functions in streamlined ultra-small and other groundwater prokaryotes. The length of transferred DNA was 10,324 ± 9,397 bp on average for the 1,046 detected GIs (10.8 Mbp total in all groundwater taxa) and 899 ± 562 bp for the 1,501 HT genes (1.3 Mbp overall) (Table S2). Therefore, while a third less GIs than individually transferred genes were detected in terms of number, the total length of sequences transferred was substantially greater with GIs, and broadly equivalent between Patescibacteria (9,689 bp per MAG) and other groundwater microorganisms (10,406 bp per MAG). Besides sequence length, it has been shown that recombination efficiency decreases exponentially with sequence divergence (Majewski et al., 2000; Majewski & Cohan, 1999). However, the presence of flanking regions of identity in exchangeable regions can remove most barriers to recombination. The shortest length of sequence homology necessary for efficient recombination can vary greatly depending on the organism, the recombination pathway used, and other factors (Thomas & Nielsen, 2005). In *Escherichia coli*, efficient recombination has been observed with as little as 23 bp of sequence homology (Shen & Huang, 1986).

### Impact of phylogenetic distance and direction of transfer

We assessed how phylogenetic distance between donor-recipient pairs associates with the metabolic function of genes acquired by Patescibacteria. Genetic exchange occurred over a wide range of phylogenetic distances based on average amino acid identities between genomes of groundwater microorganisms (as low as 38.5% AAI, Fig. 6a). Within the Patescibacteria phylum, genes of diverse metabolic functions were transferred regardless of phylogenetic distance (41.0-54.3% AAI). However, direction of HT events (between members of the Patescibacteria phylum, other groundwater microorganisms to Patescibacteria, and Patescibacteria to other groundwater microorganisms) appeared to have an impact on the metabolic function of acquired genes (Fig. 6b). A number of metabolic functions were found to be exclusively transferred between Patescibacteria (encoded by 12 out of 104 HT genes shared between Patescibacteria), and were involved in processes such as amino acid metabolism (serine hydroxymethyltransferase, GlyA), cell division (actin homolog, MreB), intracellular trafficking, co-enzyme metabolism (riboflavin biosynthesis) and inorganic ion metabolism (cation transport) (Fig. 6b, Table S6).

**Figure 6.**
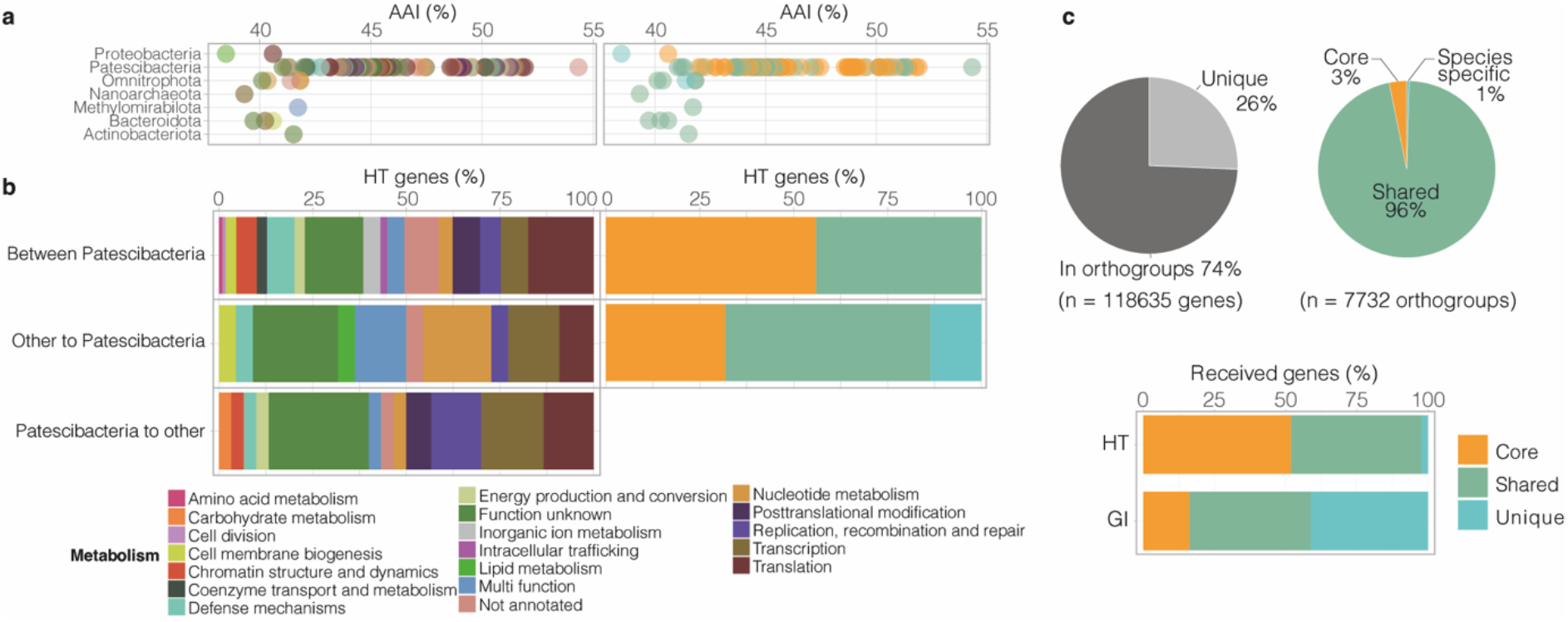
Pangenome of Patescibacteria and effect of phylogenetic distance and direction of transfer on acquired genes. (a) Impact of phylogenetic distance of HGT donor to Patescibacteria (calculated as AAI) on metabolic function of horizontally acquired genes (left) and pangenome category (right). (b) Metabolic function of HT genes according to direction of transfer. (c) Proportion of Patescibacteria total gene pool contained in orthogroups vs. unique, and breakdown of proportion of orthogroups in ‘core’, ‘shared’ and ‘species-specific’ pangenome categories (top). Proportion of horizontally acquired genes (HT genes and GI genes) by Patescibacteria taxa in each pangenome category (bottom).

Others were exclusively shared from Patescibacteria to other phyla (out of n=29 transfers overall, Fig. 6b), namely two genes associated with chromatin structure and dynamics (SWIB/MDM2 domain-containing protein) and carbohydrate metabolism (mannose-6-phosphate isomerase, COG0662). Moreover, Patescibacteria donated a greater fraction of replication, recombination and repair genes (GIY-YIG catalytic domain-containing protein, UvrD subfamily helicase, bacterial nucleoid DNA-binding protein COG0776, DNA-3-methyladenine glycosylase) to other bacteria. Among the nine HT events from Patescibacteria to DPANN archaea reported above, three were related to a glyoxalase (Table S6), which is key metalloenzyme in the glycolytic pathway involved in the detoxification of reactive methylglyoxal into D-lactate. It is a common feature across domains of life, including in Archaea (Rawat & Maupin-Furlow, 2020), and was previously reported in CPR single amplified genomes (León-Zayas et al., 2017). The reason ultra-small archaea exhibit an apparent preference for acquiring bacterial homologs is unclear.

A very small number of HGT events (two) also occurred exclusively from other prokaryotes to Patescibacteria (out of n=20 transfers overall, Fig 6b). This includes the *acpP* gene encoding an acyl carrier protein, and a gene encoding a putative lysophospholipase (COG2267, alpha beta hydrolase superfamily). Lysophospholipases catalyse phospholipid hydrolysis and generate fatty acids. A recent study coupling lipidomics and metagenomics approaches suggested that members of the Patescibacteria phylum, that lack capacities for fatty acid biosynthesis (Castelle et al., 2018), are able to recycle lipids from other bacteria for cell membrane biogenesis (Probst et al., 2020). This suggests that HGT could facilitate the sourcing of fatty acids in Patescibacteria. Phospholipase genes have also been reported in the genomes of other ultra-small CPR prokaryotes (for example, in amoeba endosymbiont *Vampirococcus lugosii*; Moreira et al., 2021). Patescibacteria were also more likely to receive genes from other groundwater microorganisms associated with nucleotide metabolism (n=2), including one thymidylate synthase genes *thyA*. The enzyme encoded catalyzes the *de novo* synthesis of dTMP (deoxythymidine monophosphate), an essential precursor for DNA biosynthesis (Voeller et al., 1995), which is a capacity already encoded by the minimal biosynthetic gene repertoire of groundwater Patescibacteria. Our results indicate that this essential function can also be horizontally acquired from other members of groundwater microbial communities.

While the number of transfer events recorded from general groundwater microorganisms to Patescibacteria and vice versa is low, limiting robust conclusions, these numbers are consistent with known associations between HGT and phylogenetic distance among prokaryotes more generally (Majewski et al., 2000). Overall, metabolic functions gained via HGT in ultra-small prokaryotes remain poorly studied, regardless of phylogenetic origin. Events identified here tend to confer Patescibacteria with critical functions, such as the ability to degrade lipids, and carry out the biosynthesis of nucleotides.

### Patescibacteria pangenome and HGT

The evolutionary fate of a HGT event is determined by fitness conferred to the recipient genome and cell, i.e., whether the acquisition of exogenous DNA is beneficial, neutral or deleterious. In order to evaluate the importance of the metabolic functions acquired through HGT in Patescibacteria, we performed a pangenome analysis of the 125 reconstructed MAGs. The pangenome of all the examined Patescibacteria contained 7,732 orthogroups, comprising 88,187 genes (74.3%) out of the total 118,635 predicted genes. Genes in the pan-genome were classified into 3 groups based on their occurrence: (1) the “core” genes, which are present in at least 2/3 of all 132 genomes, (2) the “shared” genes are found in more than one genome, but are not core genes, and (3) the “unique” genes, comprising species-specific genes (i.e. genes present in one genome only) and genes not assigned to an orthogroup. Both shared and unique genes comprise the accessory genome. Results indicated the vast majority of orthogroups (96%) were shared between the groundwater Patescibacteria, and only 3% of orthogroups (2,634 genes) comprised the core genome (Fig. 6c). This represents the lower end of core genome size among bacterial pangenomes (Bosi et al., 2017; Shin et al., 2016) and could be partially due to MAG incompleteness, but is consistent with the variation found in the core protein families by Méheust and colleagues across members of the CPR radiation (Méheust et al., 2019).

GIs brought many unique, and hence divergent, genes to the Patescibacteria pangenome (41.2% of GI genes, Fig. 6c). These observations are consistent with previous work suggesting that prokaryotes contain a higher proportion of novel genes in GIs compared with the rest of their genome (Hsiao et al., 2005). HT genes acquired by Patescibacteria species were spread across the core and the shared gene fractions of these organisms (45.8% shared and 51.9% core, Fig. 6c), and not the unique fraction. A high proportion of core or shared genes would be expected given predicted transfers occurred mostly between Patescibacteria species (83.2% of HT events, Fig. 4a). However, the lack of any unique genes reflects limitations in the detection of HT events by the MetaCHIP tool, which considers self matches to be false positive (i.e. a donor is required) (Song et al., 2019).

Streamlined genomes are characterised by low numbers of paralogs (Giovannoni et al., 2014). Accordingly, among all genes in Patescibacteria, we detected a low number of paralogous orthogroups based on the pangenome analysis results (89 genes in 36 species-specific orthogroups – i.e. 89 paralogs out of 118,635 genes). We found that no genes located on GI regions nor HT genes were paralogs of other genes in the Patescibacteria pangenome. Gene duplicates are generally under purifying selection (Lynch & Conery, 2000), and have been shown to evolve faster than non-duplicated genes with a similar level of divergence (Kondrashov et al., 2002), introducing new proteins. Nonetheless, gene duplication is usually widespread in bacterial genomes, with an estimated 7–41% of bacterial proteins encoded by paralogs (Gevers et al., 2004), and is a common strategy for creating novel gene functions for niche expansion in prokaryotes (Hooper & Berg, 2003). However, it has been quantitatively demonstrated that gene gain via HGT rather than gene duplication is the principal factor of innovation in the evolution of prokaryotes (Treangen & Rocha, 2011). Based on a comparably high proportion of HGT into Patescibacteria as for other phyla, results here demonstrate HGT, and unique genes conferred by GIs, are almost exclusively used over duplication by Patescibacteria to expand niche ranges.

### GI in Patescibacteria nzgw456 encoding a full type I restriction-modification system

Streamlined bacteria (and archaea) have been hypothesized to be largely lacking classical defense mechanisms, such as CRISPR-Cas (Clustered Regularly Interspaced Short Palindromic Repeats – CRISPR associated), restriction-modification (R-M) and toxin-antitoxin systems (Koonin et al., 2017). Similarly, the analysis of hundreds of CPR/Patescibacteria genomes recovered from diverse environments revealed that most are missing CRISPR-Cas adaptive immune systems, lost in the process of genome streamlining, and instead rely nonspecific defence, such as R-M systems (Castelle et al., 2018). Although first believed to constitute a barrier to genetic exchange between organisms (Vasu & Nagaraja, 2013), it has been recently suggested that these systems play an important role in HGT rates by increasing the frequency of exchange in organisms possessing numerous R-M systems (Oliveira et al., 2016).

Out of the 120 GIs detected in Patescibacteria we identified four encoding restriction modification systems or endonucleases in patescibacterial genomes. All four MAGs contained a high number of GIs compared to other patescibacterial counterparts (average 5 ± 2.8 SD vs 1.7 ± 1.0), and three acquired more HT per Mbp than the average for Patescibacteria overall (average 2.5 ± 1.3 SD vs 1.2 ± 1.5, Table S2). One GI in Patescibacteria nzgw456 (Paceibacteria class) encodes a full type I R-M system (Fig. 7). Type I R-M systems, which are the most complex of all four known R-M systems (Murray, 2000), comprise three subunits, namely HsdR and HsdM (required for methyltransferase activity), and HsdS (required for restriction). Co-located with genes encoding these subunits on the GI was one putative ATP-dependent endonuclease. Type I R-M related genes were also found in a GI present in Patescibacteria nzgw350, and other restriction endonucleases in nzgw410 and nzgw504 (Table S7). Acquisition of type I R-M genes via HGT was recently described in CPR bacterium *V. lugosii* (Moreira et al., 2021), however not as syntenic genes. It has been suggested that R-M systems could be used by putative epibionts as a way to source of nucleotides by degrading exogenous DNA, including that derived from viruses (Burstein et al., 2016), in addition to protecting against phage infection (Stern & Sorek, 2011).

**Figure 7.**
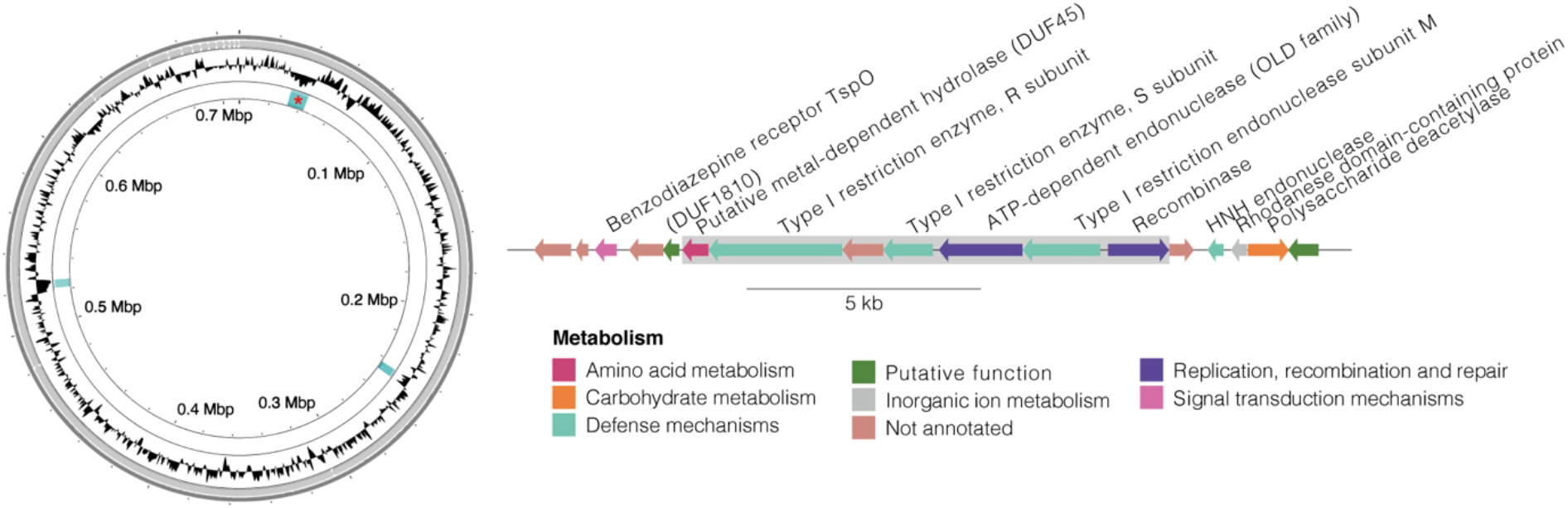
Genomic island encoding a full type I restriction-modification system in Patescibacteria nzgw456. Left: circular genome plot with following rings (inner to outer): GIs (in blue; star indicates location of GI depicted in the right panel); GC content; contig boundaries. Right: Linear map of GI (indicated by grey box) and protein-coding genes. Manual examination of GI was undertaken to further confirm boundaries using the DarkHorse prediction tool (Podell et al., 2008).

## CONCLUSION

Recent phylogenetic analyses have shed light on the massive bacterial group that is the Patescibacteria phylum/CPR (Hug et al., 2016). Due to the reductive evolution of Patescibacteria, it is essential to consider the processes of gene gain and loss in respect of the predicted symbiotic lifestyles of these ultra-small microorganisms. Results here indicate that Patescibacteria are involved in dynamic genetic exchange with their ultra-small counterparts and other members of the microbial community. We found a total 124 HT events involving Patescibacteria, 24 prophage and over 100 GIs in Patescibacteria genomes. While most transfers occurred among Patescibacteria, HGT origins were diverse and widespread across both bacterial and archaeal domains, in particular with members of the Omnitrophota and Bacteroidota phyla. A putative host symbiont pair was identified based on HGT evidence and co-occurrence across groundwater samples. Functions encoded by horizontally acquired regions comprised diverse pathways, but mainly transcription, translation and DNA replication, recombination and repair, such as ribosomal proteins and transcription factors. Findings show that acquired genes were distributed across core and accessory portions of Patescibacteria genomes, and that GIs generally introduced phylogenetically distant sequences. Some of the metabolic functions horizontally acquired by Patescibacteria are distinct from those acquired by the general groundwater communities, suggesting unique pressures on gene gain/loss dynamics occurring in ultra-small prokaryotes.

## ACKNOWLEDGMENTS

Funding was provided by a MBIE Smart Ideas grant awarded to K.M.H. (project UOAX1720). We thank D. Waite and J.S. Boey for help with bioinformatics. Computational resources were provided by New Zealand eScience Infrastructure.

The authors declare no conflict of interest.

